## Supplementary materials for "Temporal dedifferentiation of neural states with age during naturalistic viewing"

**Table S-I** Age group descriptives

| Group | *M* age | min | max | *M* qualification[1] | sex (N men) |
|---|---|---|---|---|---|
| 1 | 19.75 | 18 | 23 | 3.13 | 6 |
| 2 | 24.24 | 23 | 25 | 3.35 | 4 |
| 3 | 26.71 | 26 | 28 | 3.71 | 6 |
| 4 | 28.47 | 28 | 29 | 3.94 | 7 |
| 5 | 30.59 | 29 | 32 | 3.59 | 9 |
| 6 | 32.65 | 32 | 34 | 3.65 | 5 |
| 7 | 34.41 | 34 | 35 | 3.71 | 8 |
| 8 | 36.12 | 35 | 37 | 3.82 | 12 |
| 9 | 37.53 | 37 | 39 | 3.81 | 13 |
| 10 | 39.59 | 39 | 40 | 3.82 | 10 |
| 11 | 41.12 | 40 | 42 | 3.47 | 9 |
| 12 | 43.24 | 42 | 44 | 3.65 | 6 |
| 13 | 45.29 | 44 | 46 | 3.24 | 10 |
| 14 | 46.76 | 46 | 47 | 3.47 | 7 |
| 15 | 48.12 | 47 | 49 | 3.59 | 6 |
| 16 | 50.06 | 49 | 51 | 3.71 | 8 |
| 17 | 51.88 | 51 | 53 | 3.41 | 7 |
| 18 | 53.88 | 53 | 55 | 3.29 | 8 |
| 19 | 55.71 | 55 | 57 | 3.18 | 8 |
| 20 | 57.71 | 57 | 59 | 3.65 | 7 |
| 21 | 59.71 | 59 | 60 | 3.18 | 9 |
| 22 | 61.59 | 61 | 63 | 3.35 | 6 |
| 23 | 63.47 | 63 | 64 | 3.47 | 5 |
| 24 | 65.35 | 64 | 66 | 3.53 | 11 |
| 25 | 67.41 | 66 | 68 | 3.35 | 12 |
| 26 | 69.00 | 68 | 70 | 2.63 | 12 |
| 27 | 70.82 | 70 | 72 | 3.29 | 13 |
| 28 | 72.71 | 72 | 74 | 3.06 | 9 |
| 29 | 75.47 | 74 | 76 | 3.06 | 9 |
| 30 | 77.65 | 77 | 78 | 3.00 | 7 |
| 31 | 78.94 | 78 | 79 | 3.29 | 8 |
| 32 | 80.18 | 80 | 81 | 3.53 | 10 |
| 33 | 82.12 | 81 | 83 | 2.71 | 9 |
| 34 | 85.18 | 83 | 88 | 2.88 | 7 |

Note: [1]qualification measured in four levels; 1 = none > 16, 2 = GCSE grade, 3 = A' levels, 4 = university

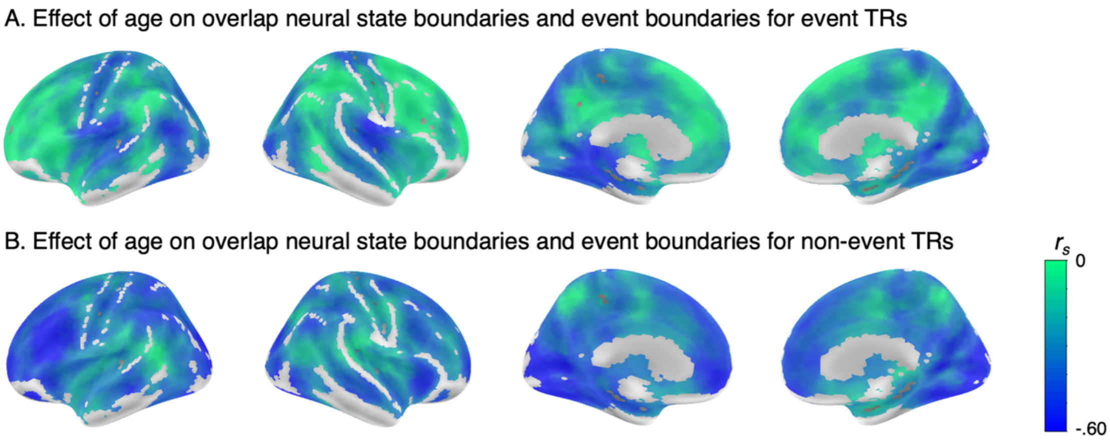


**Figure S-II** The effect of age on boundary occurrence for TRs that overlap with perceived events (A) and TRs that do not (B).

Figure S-II visualizes the difference between alignment and non-alignment with event boundaries for the correlation between age and neural boundary occurrence. In both event and non-event TRs we see an overall decrease in boundary occurrence with age (i.e., longer states), however this effect is stronger for non-event TRs (mean correlation across all searchlights $r_{s} =$ -0.36) than event TRs ($r_{s} =$ -0.21). This explains why we observe longer neural states with increasing age without a decrease in overlap between event and neural state boundaries.

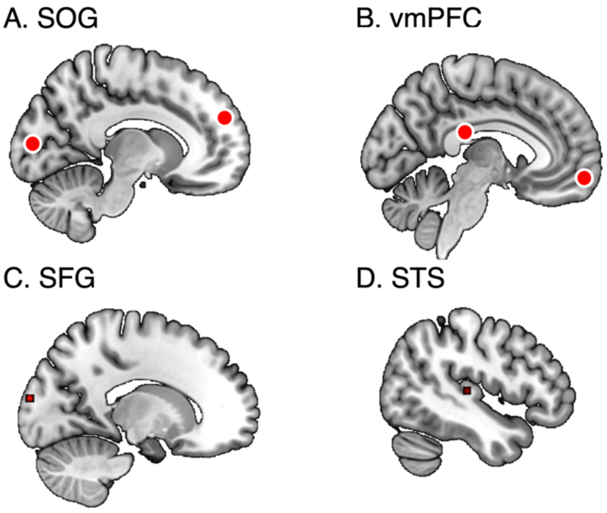


**Figure S-III** Location of selected searchlights for single subject GSBS and simulations. Top row: searchlights with a strong effect of age on neural state duration. A. Superior occipital gyrus, SOG, -15 x -96 x 14, SL 1874; B. vmPFC, -6 x 57 x -13, SL 2466. Bottom row: searchlights with a high overlap between perceived event boundaries and neural state boundaries. C. Superior frontal gyrus, SFG, -9 x 51 x 26, SL 2463; D. superior temporal gyrus, STS, -45 x -24 x 6, SL 692. Coordinates in MNI space.

Figure S-III visualizes the location of the four searchlights that are used in searchlight specific analyses. The two searchlights in the top row are also visualised in the main manuscript in the section *Increase in neural state duration with age*, figure 3. Those are the searchlights with the highest correlation between age and median state duration. The searchlights on the bottom row are those with a high overlap between perceived event boundaries and neural state boundaries. All four were used for single-subject GSBS to investigate the overlap between neural states and events on a single-subject level.

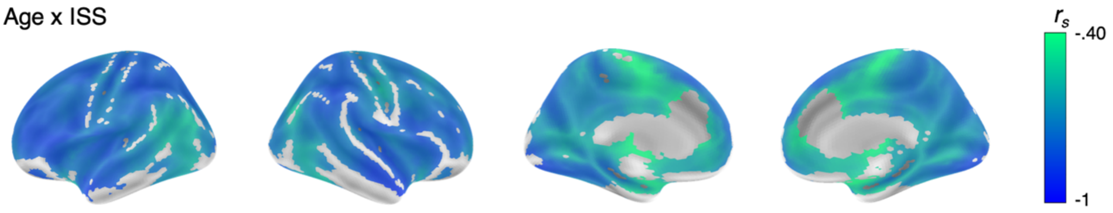


**Figure S-IV** The effect of age on ISS. Age has a negative effect on ISS across the cortex indicating that with increasing age the neural signal is less similar.

Figure S-IV shows the effect of age on intersubject synchrony (ISS). Age had the strongest negative effect on ISS bilaterally in the temporal pole, the dorsolateral PFC, and the inferior parietal lobe. With increasing age the neural signal is less similar. This spatial pattern of correlations only had a small overlap with regions where the effect of age on neural state duration was strongest ($r_s$ = -.09, $p$ < .001, figure 2 main manuscript), indicating that age affects ISS and state duration largely in different regions of the brain. This suggests that there is a different driving force behind the age-related differences in neural state duration and the differences in ISS.

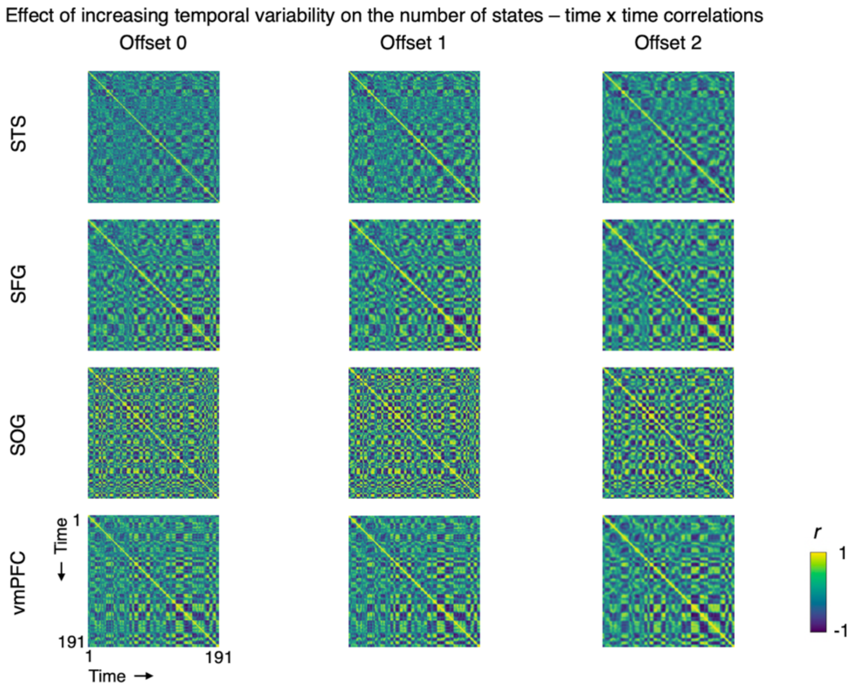


**Figure S-V** The effect of increasing temporal variability with 1 or 2 TRs relative to the data of the youngest subgroup, visualized in time-by-time matrices for four selected searchlights, two that show a strong effect of overlap (top rows) and two with strong effect of age on duration (bottom rows).

Figures S-V till S-VII show results of simulations. S-V visualises the how interindividual temporal variability in the occurrence of state boundaries affected the duration of neural states by shifting neural state boundaries of the youngest group by one or two TRs. As can be seen by comparing figure S-V with figure S-VIII, the time-by-time matrices based on simulations look dissimilar from those of older adults.

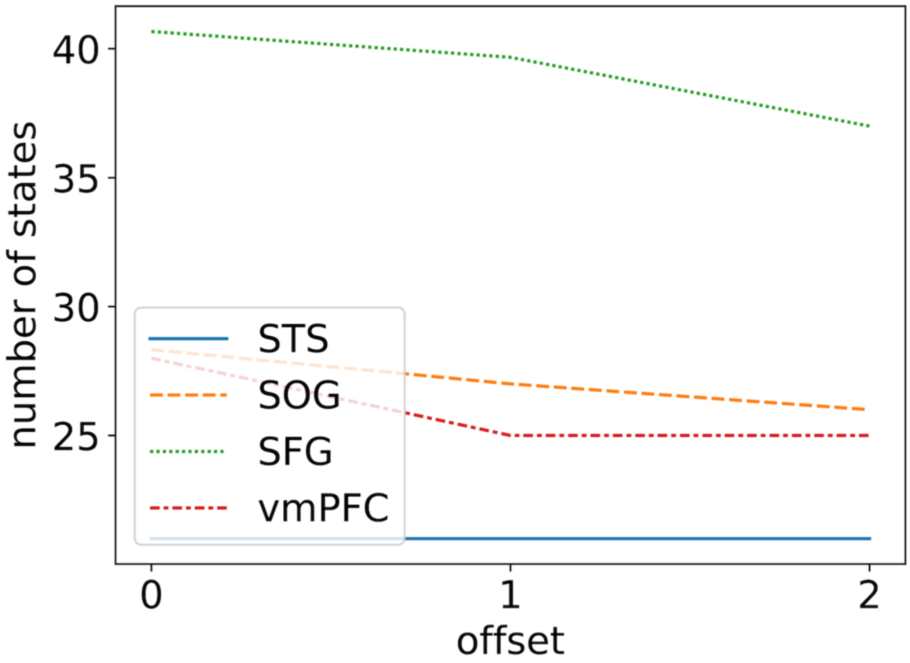


**Figure S-VI** The effect of increasing temporal variability on the number of states for four selected searchlights.

Figure S-VI shows that the number of states tend to decrease slightly with a simulated increase in inter-individual variability. Importantly, these effects are not in the range that we observe with aging. While with advancing age, the number of states can decline very steeply (e.g. from 41 to 25 states in the SOG), the simulations show a much smaller decrease (largest difference is from 41 to 37), suggesting that increased variability in the timing of state boundaries cannot explain the observed increase in neural state durations with age.

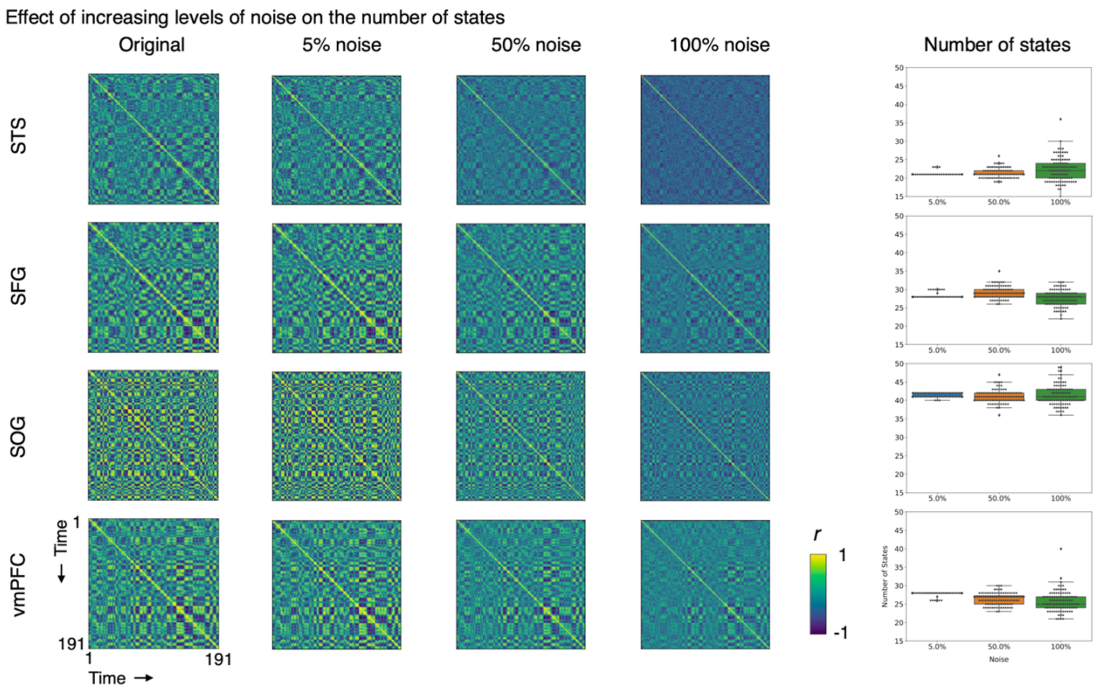


**Figure S-VII** The effect of increasing random noise relative to the data of the youngest group, visualized in time-by-time matrices for four selected searchlights.

In the next simulation, we investigated whether decreased signal-to-noise levels could explain the observed increase in neural state durations with age. Figure S-VII visualises the effect of adding increasing levels of random noise to the data of the youngest group to investigate if this could explain the increase in neural state durations with age. While visually this made the matrices look more similar to those of older adults (see figure S-VIII), we did not observe a systematic decrease in the number of neural states with higher levels of noise.

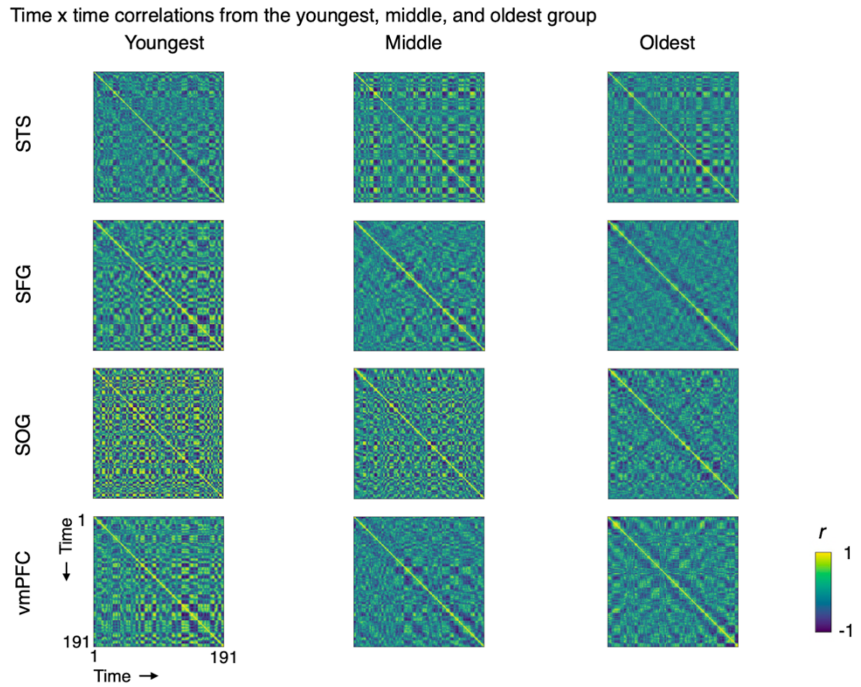


**Figure S-VIII** Time-by-time matrices from the youngest, middle, and oldest group for four selected searchlights as reference for effects of age.

Figure S-VIII shows the time-by-time matrices for the four selected searchlights for the youngest, middle, and oldest group. These are based on the actual data and can be used as a reference for the simulation results.

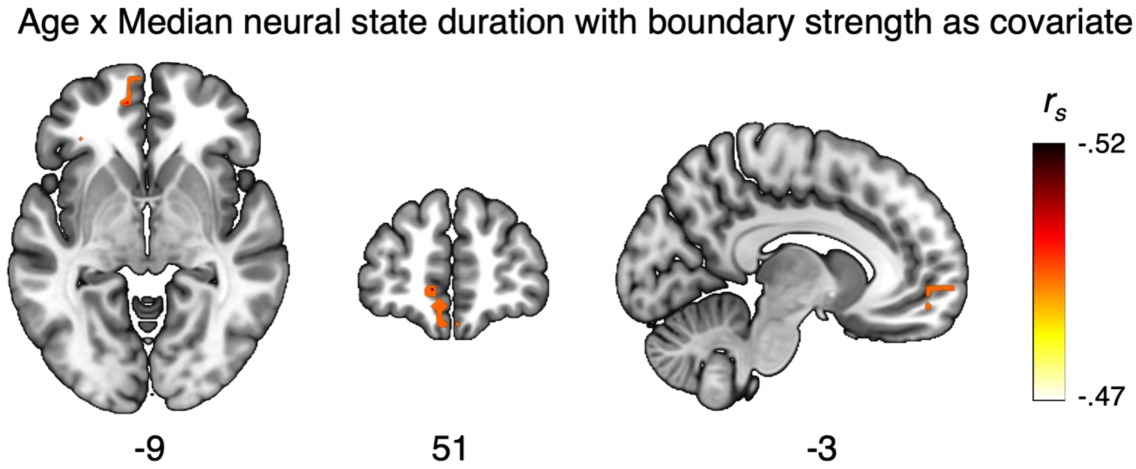


**Figure S-IX** The correlation between age and state duration with boundary strength as covariate.

Our results show that neural state boundaries were weaker in older adults which might explain part of the observed lengthening of neural states with age. Figure S-IX shows that the effect of age on state duration persisted in the vmPFC whilst taking boundary strength into account. This suggests that the lengthening of neural states cannot be fully explained by the weakening of neural states with age.

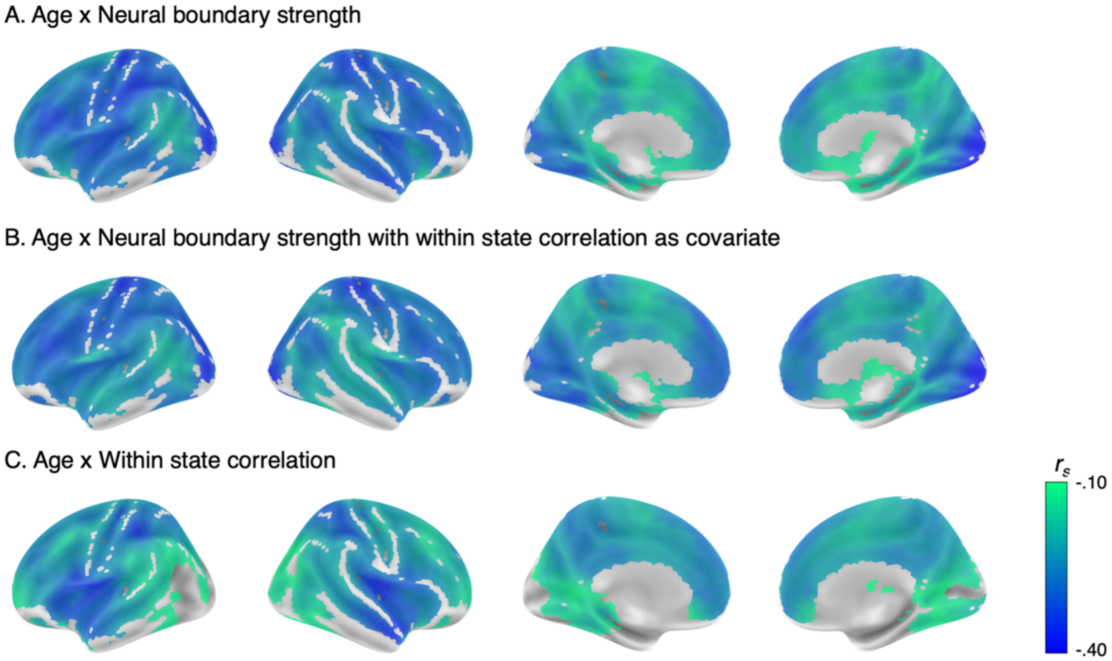


**Figure S-X** The effect of age on neural state boundary strength and within state correlation. A) The effect of age on boundary strength, a replication of the original analysis reported in the section *Weaker neural state boundaries only partly explain the effect of age on state duration,* figure 7. B) The effect of age on boundary strength adjusted for within state correlations. C) Correlation between within state correlations and age.

Weaker boundaries might result from decreased similarity within neural states (resulting in a less coherent neural pattern per state or more noisy data) as well as from decreased distinction of neural patterns across states. Therefore, we also tested whether the correlation between age and boundary strength remained after correcting for average within state correlation (as reduced within-state coherence could give the impression that boundary strength decreases with age). However, the effect of age on boundary strength remained, suggesting that weaker state boundaries with age cannot be explain by a lack of coherence within states. Although it should be noted that within-state correlations also declined with age.
